# Improving xylose consumption in *Rhodotorula toruloides* through heterologous expression of xylose reductase and xylulokinase

**DOI:** 10.1101/2023.05.10.540254

**Authors:** Paola Monteiro de Oliveira, Marina Julio Pinheiro, Juliano Sabedotti De Biaggi, Artjom Tšitšerin, Eliise Tammekivi, Koit Herodes, Nemailla Bonturi, Petri-Jaan Lahtvee

## Abstract

The oleaginous yeast *Rhodotorula toruloides* is a promising host for sustainable bioproduction due to its capacity to naturally utilize xylose present in lignocellulosic biomass, an abundant and renewable resource. However, its xylose consumption pathway is still not completely understood. To better understand the potential limitations in xylose utilization in *R. toruloides*, heterologous xylose reductase from *Scheffersomyces stipitis*, together with the native and heterologous xylulokinases from three different microorganisms (*Scheffersomyces stipitis, Candida intermedia*, and *Escherichia coli*) were overexpressed solely and in combination. The overexpression of xylulokinases showed more significant improvements in terms of xylose consumption rate compared to the single overexpression of xylose reductase. When the heterologous xylulokinase from *Escherichia coli* was overexpressed, the specific xylose consumption rate was improved by 66% and the maximum specific growth rate by 30% compared to the parental strain. The xylose specific consumption rate increased by 146% and the maximum specific growth rate increased by 118% when heterologous genes for xylose reductase and xylulokinase from *E. coli* were overexpressed together. These results suggest that the low expression of xylulokinase in *R. toruloides*, which has been reported previously, could limit its sugar consumption, while supporting higher lipid accumulation in this yeast.

## 1. Introduction

Plant biomass is a renewable resource and one of the most abundant carbon reserves on Earth. It can be used as a feedstock to produce chemicals, energy, and other valuable products for a bio-based economy. This type of economy has generated interest over the years, aiming for independence from fossil fuels, and a reduction of waste and greenhouse gas emissions [1]. Plant cell walls are composed of lignocellulose, a polymer made of cellulose, hemicellulose, and lignin [2,3]. Hemicellulose represents 14-40% of the lignocellulose and is the second most abundant polysaccharide found in nature, being a heterogeneous polymer made of pentoses (xylose being the most abundant), hexoses, and acetic acid [2,4]. The carbohydrates present in lignocellulosic biomass need to be hydrolysed first into fermentable sugars to be available for conversion into bioproducts by a wider range of microorganisms [3]. When this process occurs, the resulting mixture is called linocellulosic hydrolysate. Due to its high availability, xylose utilization in an efficient way is essential for developing bio-based economy [5].

Microbial cell factories (MCF) are microorganisms able to convert substrates, including lignocellulosic hydrolysates, into bioproducts [6]. *Saccharomyces cerevisiae*, the most studied eukaryote microorganism, is widely used as a MCF for different biotechnological goals due to its well-established fermentation process in large-scale production [7]. Besides that, since the genome of this yeast was completely sequenced by Goffeau et al. (1996) [8], a range of genetic tools were developed, turning it into a widely used model organism for metabolic engineering and synthetic biology applications [9]. Although, this yeast is not able to consume xylose naturally, genetic modifications and laboratory evolution have been used to make this strain able to grow and efficiently metabolise also pentose sugars [10].

Xylose uptake in yeasts usually happens in two steps. In the first step, D-xylose is reduced to xylitol by xylulose reductase (XR) in an oxidoreductive pathway (XR-XDH) (Figure 1). In the second step, xylitol is reduced to D-xylulose (Figure 1) by xylitol dehydrogenase (XDH). Then, D-xylulose is phosphorylated to D-xylulose-5-phosphate by xylulokinase (XK), and D-xylulose-5-phosphate is further metabolized in the pentose phosphate pathway (PPP). In bacteria, the most common xylose consumption pathway is called the xylose isomerase pathway (XI) in which D-xylose is isomerized to D-xylulose in a reaction catalysed by xylose isomerase (XI) in one step (Figure 1) [11–13]. Moreover, some bacteria can metabolize xylose through the Weimberg and Dahms pathways, which are non-phosphorylative (Figure 1)[14]. In these pathways, xylose can be converted into α-ketoglutarate (Weimberg pathway) or pyruvate and glycolaldehyde (Dahms pathway) through a 2-keto-3-deoxy-xylonate intermediate [14,15].

**Figure 1.**
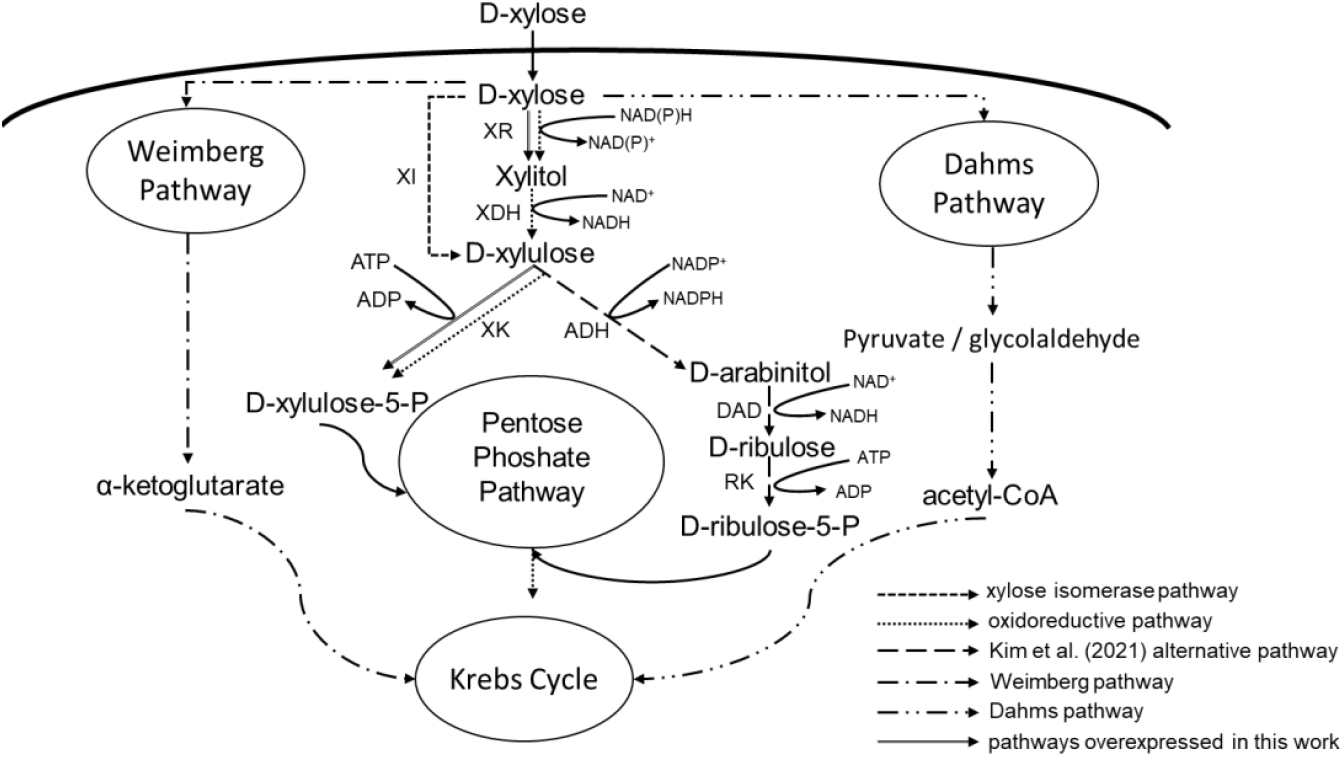
Xylose uptake metabolisms. XI: Xylose isomerase; XR: Xylose reductase; XDH: Xylitol dehydrogenase; XK: Xylulokinase.

*S. stipitis* is a MCF well-known for its to natural ability to convert xylose with high efficiency using the XR-XDH pathway [15–17]. This strain is studied for ethanol production from hydrolysates, but its genome editing tools are not as well-known or studied as those of *S. cerevisiae* [18,19]. However, XR and XDH from this yeast (ssXR and ssXDH, respectively) have been widely used in metabolic engineering of the xylose consumption pathway in various organisms [10,17,20], as the ssXR is able to use both NADH and NADPH as co-factors [21–23]. This cofactor versatility allows the yeast to adapt to the redox demands in response to the environment. For example, it was shown an inverse relation between oxygen supply and NADH-dependence of *S. stipitis* XR. This happens because, when oxygenation is high, more NADH is needed by the electron transfer chain, so it is preferable that XR used NADPH instead [25]. Most of xylose-consuming yeasts reduce D-xylose to xylitol catalysed by an NADPH-dependent XR, leading to an unbalance of cofactors once XR uses NADPH and XDH produces NADH (Figure 1).

Another MCF that has been studied as a potential biotechnological microorganism and has a native xylose consumption pathway is *Rhodotorula toruloides* [26]. It is an non-conventional and oleaginous yeast which is classified as basidiomycete [27]. Furthermore, de Oliveira et al. (2021) [28] showed that this strain was able to grow in hemicellulosic hydrolysate in high concentration showing the shortest lag phase (by consuming all sugars in 30 h of cultivation) when compared to other yeast cell factories, including *S. stipitis*. Besides being able to consume different carbon sources, *R. toruloides* also has a high tolerance for inhibitory compounds in lignocellulose hydrolysate, being considered a potential MCF for biorefineries [26]. Pinheiro et al. (2020) [29] observed that XK was not detected in their proteomic analysis of *R. toruloides* CCT 7815 and suggested that this fact could limit its xylose catabolism in this yeast. Kim et al. (2021) [30] also observed very low XK expression in transcriptomic analysis of *R. toruloides* IFO0880 genome and suggested an alternative xylose pathway where D-xylulose is converted to D-arabinitol, forming D-ribulose and then ribulose-5-phosphate instead of forming D-xylulose-5-phosphate from xylulose (Figure 1). A similar pathway was also suggested for *R. toruloides* CCT 7815 strain based on an enzyme constrained model [31], where regeneration of NADPH was also identified as one of the potential bottlenecks in the *R. toruloides* growth in xylose.

The aim of this work was not only to better understand the xylose utilization pathway in *R. toruloides* but also to improve its consumption rate and understand how the production of carotenoids and lipids are affected by it. The overexpression strains of: (1) a heterologous XR; and (2) native or (3) heterologous XK, were constructed and characterised individually and synergically. For this, a heterologous XR from *S. stipitis* was inserted into *R. toruloides* aiming to understand the cofactor influence on the xylose consumption in this strain. To remediate the lack of or low levels of XK shown by [14,26], we also over-expressed XK using a strong native promoter in *R. toruloides*, to study the effect of this enzyme in the xylose catabolism in this yeast.

Our results showed that the highest rates for specific-xylose consumption rate - g of xylose consumed per g of dry cell weight per hour (r_xyl_) - and maximum specific growth rate (μ_max_) were observed when heterologous XR and XK were overexpressed together (r_xyl_ = 0.44 g_xyl_.g_dcw_^-1^.h^-1^ and μ_max_ = 0.13 h^-1^, respectively). The strain grew 118% faster and consumed xylose 146% faster when compared with the parental strain and there no significant difference was observed in carotenoid production and lipid content. Between the XK overexpression (homologous and heterologous), the best result was achieved when XK from *Escherichia coli* was overexpressed (r_xyl_ = 0.30 g_xyl_.g_dcw_^-1^.h^-1^ and μ_max_ = 0.08 h^-1^), showing an improvements of 66% in specific-xylose consumption rate and 30% in maximum specific growth rate when compared with the parental strain.

## 2. Materials and Methods

### 2.1. Microorganisms and culture media

The strain used in this work was *Rhodotorula toruloides* SBY92 [32] – called parental strain here – a strain derived from *R. toruloides* CCT7815 (Fundação André Tosello, Campinas, Brazil) but with an extra copy of its native carotenoids biosynthesis genes. All the strains used in this study are shown in Table 1. Competent *Escherichia coli* DH5α grown in LB medium (10 g.L^-1^ tryptone, 5 g.L^-1^ yeast extract, 10 g.L^-1^ NaCl) supplemented with chloramphenicol (25 mg.L^-1^) or kanamycin (30 mg.L^-1^) at 37 °C was used for multiplication and storage of all plasmid constructions. For yeast transformation, the cells were cultivated in YPD medium (yeast extract, 10 g.L^-1^; peptone, 20 g.L^-1^; and glucose, 20 g.L^-1^). The *R. toruloides* inoculum was prepared in YPX medium (yeast extract, 10 g.L^-1^; peptone, 20 g.L^-1^; and xylose, 20 g.L^-1^). The yeast was cultivated in buffered mineral media (ammonium sulphate, 0.83 g.L^-1^; magnesium sulphate, 0.25 g.L^-1^, monopotassium phosphate, 3 g.L^-1^; dipotassium phosphate, 5.25 g.L^-1^; vitamins and trace elements solutions [33] supplemented with xylose at defined concentration.

**Table 1.** Transformed strains studied on this work. ssXR: heterologous xylose reductase from *S. stipitis*; rtXK native xylulokinase; ecXK: heterologous xylulokinase from *E. coli*; ciXK: heterologous xylulokinase from *C. intermedia*; ssXK: heterologous xylulokinase from *S. stipitis*. pXYL: promoter from xylose reductase (XYL1,); pADH2: promoter from alcohol dehydrogenase 2 (ADH2,); tNOS: terminator from *Agrobacterium tumefaciens* nopaline synthase (tNOS,); HYG and NAT: resistance gene for hydromycin and nourseothricin, respectively. PEX10 and PEX11: Peroxisomal Biogenesis Factors 10 and 11.

| Strain | Construction | Reference |
| --- | --- | --- |
| SBY92 | $\Delta ku70::$ pXYL-G418-tNOS-pXYL-crtE-tNOS-pGPD-crtI-tNOS-pADH2-crtYB-tNOS | Bonturi et al.; 2022 |
| SBY137* | $\Delta pex10::$ NAT-pXYL-ssXR-tNOS | This study |
| SBY138* | $\Delta pex10::$ NAT-pXYL-rtXK-tNOS | This study |
| SBY139* | $\Delta pex10::$ NAT-pXYL-ssXK-tNOS | This study |
| SBY140* | $\Delta pex10::$ NAT-pXYL-ecXK-tNOS | This study |
| SBY141* | $\Delta pex10::$ NAT-pXYL-ciXK-tNOS | This study |
| SBY165** | $\Delta pex11::$ HYG-pADH2-ssXR-tNOS | This study |
| SBY166*** | $\Delta pex11::$ HYG-pADH2-ssXR-tNOS | This study |

\* SBY92 as background. \*\* SBY140 as background. \*\*\* SBY141 as background.

### 2.2. Construction of expression cassettes of XR and XK and transformation into yeast strains microorganisms and culture media

The expression cassette constructions were done using the recently developed Golden Gate Assembly platform for *R. toruloides* – RtGGA [32]. All parts were designed to bear pre-defined 4-nt overhangs and BsaI recognition sites in both ends and cloned in TOPO vector (TOPO™ XL-2 Complete PCR Cloning Kit, Thermo Fischer). The genomic DNA was extracted according to Lõoke et al. (2011) [34] for the cassette assembly by polymerase chain reaction (PCR) using Platinum SuperFi Master Mix (Thermo Fisher Scientific, Lithuania) following the manufacturer’s instruction for high GC content-templates. After the reaction, the amplicons were separated in a 1% agarose gel after electrophoresis at 140 V and identified by size based on gene ruler Plus DNA Ladder (Thermo Fisher Scientific, Lithuania). Amplicons were purified using Favorprep™ GEL/PCR Purification Kit (Favorgen, Austria) and quantified by Nanodrop (Thermo Fisher, Whaltan, USA). All primer’s sequences used in this study can be found in Supplementary Table S1. The sequence for xylose reductase from *Scheffersomyces stipitis* CBS 6054 (ssXR, XM_001385144), and the xylulokinases from *E. coli* (ecXK, Gene ID: 948133), *S. stipitis* CBS 6054 (ssXK, XM_001387288) and *Candida intermedia* CBS 141442 (ciXK, SGZ54087) were codon optimized for expression in *R. toruloides* using Benchling (https://benchling.com) and the genes were synthetized by Twist Biosciences (San Francisco, USA). The native gene of xylulokinase (rtXK, RHTO_04556) was PCR-amplified from *R. toruloides* CCT7815 genomic DNA. The insertional regions for the cassette genomic integration consisted in five hundred base pair fragments upstream (insUP) and downstream (insD) of the Peroxisomal Biogenesis Factors 10 (PEX10, RHTO_05319) and 11 (PEX11, RHTO_03603). The aforementioned sequences were amplified from the genome.

The promoters from the genes alcohol dehydrogenase 2 (ADH2, *Addgene ID* 195015), xylose reductase (XYL1, *Addgene ID* 195016), terminator from *Agrobacterium tumefaciens* nopaline synthase (tNOS, *Addgene ID* 194997) and markers, consisting in the resistance genes expression cassettes for hygromycin (*HYG, Addgene ID 195013*) or nourseothricin (*NAT, Addgene ID 195014*), were obtained from the library created by Bonturi et al. (2022) [32] and deposited at Addgene (https://www.addgene.org/).

For assembly of the expression cassettes, a modified pGGA vector (New England Biolabs, USA) containing a red fluorescent protein and carrying chloramphenicol resistance gene was used for facilitating the screening of positive bacterial transformants [32]. The GGA reaction was carried out as following: 75 ng of modified pGGA; each insert added to a mass corresponding to 2:1 molar ratio (insert:vector); 2 μL of T4 DNA Ligase Buffer, 1 μL of T7 DNA ligase, 1 μL of BsaI, and nuclease-free H_2_O up to 20 μL (New England Biolabs, USA). The reaction was performed in 30 cycles of 37 °C for 5 min followed by 16 °C for 5 min and finalized by incubation at 60 °C for 5 min. Subsequently, the reaction mixture was used for *E. coli* DHα5 transformation. White colonies were screened for investigation of the corrected assembly. The plasmids were recovered using the Favorprep™ Plasmid DNA Extraction Mini Kit (Favorgen, Wien, Austria) and the assemblies were confirmed by PCR and sequencing.

The expression cassette was PCR-amplified from the plasmid and transformed into *R. toruloides* SBY92 or SBY140 using lithium acetate method described by Nora et al. (2019) [35]. The cells were plated in YPD agar plates, containing either 100 μg.mL^-1^ of (NAT) or 50 μg.mL^-1^ of HYG and incubated at 30 °C for at least two days. The transformants were initially screened for cell growth in xylose medium using microplate cultivation in terms of maximum specific growth rate (μ_max_) and length of the lag phase, followed by more detailed characterization using shake flasks.

### 2.3. Initial growth screening and characterization of the transformed strains with XR and XK

The initial screening of transformant strains was done in a 96-well microplate with 150 μL of mineral medium with vitamins and minerals and 1 g.L^-1^ of xylose. The cultivation was carried out at 30 °C in constant mixing on BioTek Synergy H1 Multi-Mode Reader (Agilent Technologies Inc., Santa Clara, CA, USA) for 72 h. The online OD600 measurements were done every 30 min for growth estimations. All cultivations were performed in at least five replicates. The transformants that showed best performance in terms of μ_max_ and length of lag phase were chosen for further characterization.

The characterization of selected strains was done in 250 mL-flasks using 50 mL of mineral medium plus vitamins and minerals with 30 g.L^-1^ of xylose, resulting in a C/N ratio of 80 (mol.mol^-1^). The cultivation started with an OD600 of 0.5 with cells from the inoculum. The growth profile was analysed through OD600 measurement during the experiment. For xylose analysis, the samples were taken and centrifuged to remove the cells and kept at -20 °C until analysis. All cultivations were performed in triplicate. Statistical analyses were done using Microsoft Excel (2016) functions FTEST and T.TEST to, respectively, analyse the difference between sample’s variances and means (p-value ≤ 0.05).

### 2.4. Analytical methods

Cell growth was measured by OD600 nm using Ultrospec 10 Cell Density Meter (Biochrom US, Holliston, Massachusetts, USA). The concentration of the dry cell biomass (g_dcw_.L^-1^) was determined gravimetrically by collecting the broth and harvesting the cells by centrifugation at 4000 *g* for 10 min, followed by washing the pellet twice with purified water and freezing at -80 °C. Then the cells were lyophilized (Scanvac CoolSafe 110-4, Labogene, Denmark) and weighted. This material was also used for further carotenoids and lipids extraction and quantification. Xylose, arabitol and xylitol were quantified by HPLC with a RID-20A detector (LC-2050C, Shimadzu, Kyoto, Japan). The column used in the analyses was Rezex RPM Monosaccharide Pb+ 300 × 7.8 mm (Phenomenex, Torrance, United States) at 85 °C. Purified water was used as mobile phase at a flow rate of 0.6 mL.min^-1^. The carotenoid extraction was carried out according to Pinheiro et al. (2020) [29]. The calibration curve was made using β-carotene (Alfa Aesar, MA, USA) and the quantification was done by measuring absorbance at 448 nm [36] with a spectrophotometer. For the lipid quantification and determination of the lipid profile, direct transesterification of the dry biomass [37] was used. This derivatization produced fatty acid methyl esters that were thereafter quantified using gas chromatography–mass spectrometry (GC-MS) [38]. An Agilent (Santa Clara, CA, USA) 7890A GC instrument connected to an Agilent 5975C inert XL mass selective detector (MSD) with a triple-axis detector and an Agilent G4513A autosampler were used. An Agilent DB-225ms capillary column (30 m x 0.25 mm diameter, 0.25 μm film thickness) with a (50%-cyanopropylphenyl)-methylpolysiloxane stationary phase was selected because of its efficient separation of fatty acid methyl esters.

## 3. Results

### 3.1. Construction of an improved xylose consumption pathway in R. toruloides

In order to improve the *R. toruloides* metabolism in xylose cultivation, three strategies were tested. First, a heterologous XR from *S. stipitis* (ssXR) was inserted into *R. toruloides* genome, since this gene is able to use two different cofactors, NADPH or NADH [21– 23,34,35]. This characteristic could favour lipid accumulation in *R. toruloides* by increasing the NADPH pool. According Pinheiro et al. (2020) [29], genome-scale modelling showed that the flux through rtXR is the main NADPH sink in *R. toruloides* metabolism in xylose cultivation. Using the ssXR, the NADPH oxidation could be suppressed by the use of NADH instead. Secondly, a native (rtXK) or heterologous XKs (ssXK, ciXK, and ecXK) were overexpressed, as the rtXK showed a low expression levels in transcriptomic and proteomic data when *R. toruloides* was cultured in xylose [14,26]. Thirdly, the two previous strategies were merged to test the combination of the applied changes. In order to achieve that, the ssXR was chosen, and among the options XK, ecXK (which demonstrated superior performance compared to the other XK options), and ciXK were selected.

The genes XR and XK from *S. stipitis* were used in this work since both enzymes are commonly used to enhance xylose metabolism in microorganisms [10]. XK from *C. intermedia* was also selected since this strain has a high capacity for xylose conversion and is one of the fastest when growing in the presence of this sugar, with a μ_max_ similar to when it is grown in glucose (0.50 h^-1^ and 0.49 h^-1^, respectively [36–38]).

A strong constitutive promoter, pXYL, was used for all the constructions in this study [32]. In case of the single transformants, the selected XR and XK genes were integrated in the locus of PEX10 (Peroxisomal Biogenesis Factors 10). Deletion of PEX10 is a commonly used strategy to improve lipid accumulation in *Y. lipolytica* by disrupting the β-oxidation pathway [39,40]. Despite that, Zhang et al. (2016) [46] observed that when PEX10 was deleted in *R. toruloides*, the strain grew slower and lipids production decreased. Furthermore, Schultz et al. (2022) [47] did not manage to obtain any transformant when deleting PEX10 in *R. toruloides*, but *R. toruloides* IFO0880, a haploid strain, was used in this study. In our study, we utilized *R. toruloides* CCT7815 [32], polyploidy yeast, which likely enabled the successful deletion of PEX10. In previous experiments in our work group, we did not observe any changes when PEX10 was deleted, suggesting that it could be a neutral site for integration for SBY92. The cassettes were efficiently assembled using the rtGGA platform and transformed into the yeast *R. toruloides* SBY92 (Table 1) [32]. To assess the effect of combining the overexpression of both a heterologous XR and various XKs the rtGGA was used to construct a new cassette for ssXR using the medium strength pADH2 promoter [32], HYG marker, and PEX11 (Peroxisomal Biogenesis Factors 11) as integration site. Pinheiro et al. (2020) [29] suggested that the down-regulation of β-oxidation could partially explain the increase in lipid accumulation and showed through proteomics analysis that PEX11 was the most abundant peroxisomal biogenesis protein in *R. toruloides*. Therefore, the deletion of *PEX11* could be efficient in increasing the lipid accumulation. The double transformant strain was obtained by integrating the *HYG*-pADH2-*ssXR* cassette into SBY140 or SBY141 (Table 1). The obtained single and double transformants were, first, subjected to an initial screening, followed by a more detailed characterisation in shake flasks.

### 3.2. Characterization of transformed strains with heterologous expression of the gene xylose reductase from Scheffersomyces stipitis

After the yeast transformation with heterologous XR, three colonies (named A, B, C) were selected from the YPD plate for an initial screening using microplate cultivation. Based on the calculated μ_max_ and the length of the lag phase of the candidates (Supplementary Figure 1A and 2), one colony was selected for further characterization in shake flasks cultivation, from which μ_max_ (h^-1^), xylose specific consumption rate (r_xyl_: g_xyl_.g_dcw_^-1^.h^-1^), and biomass yield on xylose (Y_sx_, g_dcw_.g_xyl_^-1^) were calculated. This selected strain showed an improvement in all these parameters and was named SBY137. When strain SBY137 was compared with the parental strain, it showed an improvement of 46% in the μ_max_ (Figure 2a), 26 % in the r_xyl_ (Figure 2b), and 38% in Y_sx_ (Figure 2c), where r_xyl_ showed statically significant improvement (p value ≤ 0.05) - values in Table 2.

**Table 2.** Yields and specific rates calculated for *R. toruloides* transformed and parental strains. The asterisk (*) means that values are statistically significant different with p-value ≤ 0.05 when compared with the values shown by the parental strain.

| | $\mu_{\max} (\text{h}^{-1})$ | | $r_{\text{xyl}} (\text{g}_{\text{xyl}}.\text{g}_{\text{dcw}}^{-1}.\text{h}^{-1})$ | | $Y_{\text{sx}} (\text{g}_{\text{dcw}}.\text{g}_{\text{xyl}}^{-1})$ | |
| --- | --- | --- | --- | --- | --- | --- |
|  | Average | SD | Average | SD | Average | SD |
| SBY92 (parental strain) | 0.06 | 0.01 | 0.18 | 0.01 | 0.36 | 0.05 |
| SBY137 | 0.10 | 0.02 | 0.23* | 0.02 | 0.25* | 0.02 |
| SBY138 | 0.06 | 0.01 | 0.28* | 0.02 | 0.22* | 0.04 |
| SBY139 | 0.06 | 0.00 | 0.26* | 0.01 | 0.24* | 0.02 |
| SBY140 | 0.08* | 0.00 | 0.30* | 0.03 | 0.28 | 0.03 |
| SBY141 | 0.07 | 0.01 | 0.21* | 0.02 | 0.33 | 0.02 |
| SBY165 | 0.13* | 0.00 | 0.44* | 0.01 | 0.28 | 0.00 |
| SBY166 | 0.12* | 0.01 | 0.27* | 0.02 | 0.45* | 0.02 |

**Figure 2.**
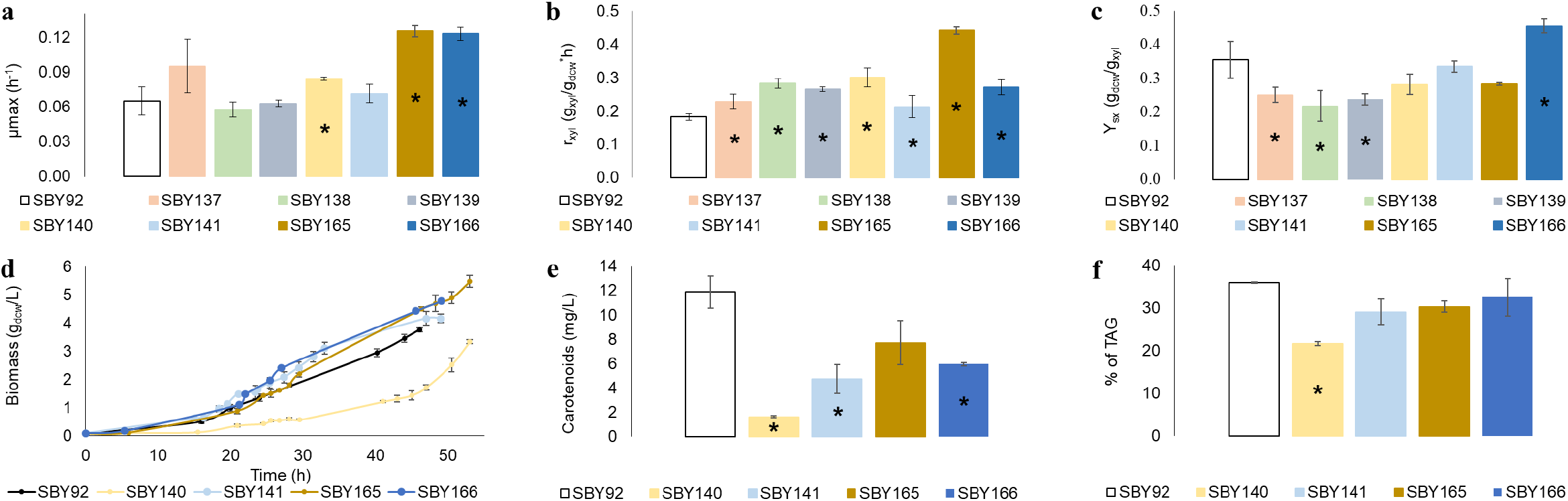
(**a**) maximum specific growth rate (μmax: h^-1^), (**b**) xylose specific consumption rate (r_xyl_: g_xyl_.g_dcw_^-1^.h^-1^), (**c**) biomass yield on xylose (Y_sx_, g_dcw_.g_xyl_^-1^), (**d**) biomass in g_dcw_.L^-1^, (**e**) total carotenoid concentration in mg.L^-1^ after 48 h of cultivation, (**f**) TAG composition in biomass in % after 48 h of cultivation. SBY92: *R. toruloides* parental strain; SBY137: transformed strain with ssXR; SBY138: transformed strain with rtXK; SBY139: transformed strain ssXK; SBY140: transformed strain with ecXK, SBY141: transformed strain with ciXK; SBY165: double-transformant strain with ssXR and ecXK; SBY166: double-transformant strain with ssXR and ciXK. The asterisk (*) means that values are statistically significant different with p-value ≤ 0.05 when compared with the values shown by the parental strain.

### 3.3. Characterization of transformed strains with native and heterologous expression of the gene xylulokinase

Next, the native XK was overexpressed in *R. toruloides*, resulting in several colonies. Based on the preliminary screening results (the three best candidate colonies shown in Supplementary Figure 1b and 2), the most promising candidate was chosen for further characterization in triplicate shake flasks (SBY138). This strain did not show a significant improvement in μ_max_ but showed a statistically significant (p-value ≤0.05) improvement of 56% in r_xyl_ (Table 2; Figure 2b) when compared to parental strain (SBY92).

Heterologous genes for XKs from three different microorganisms were also overexpressed in *R. toruloides*. Some colonies were selected for the first microplate screening. The μ_max_ were calculated for all the candidates (Supplementary Figure 2) and based on these results three strains that showed improvement in μ_max_ were selected for further characterization in shake flasks (SBY139, SBY140, and SBY141).

The same previously mentioned parameters were calculated for the three strains (Table 2; Figure 2a-c). The strain SBY140 demonstrated an improvement of 66% in r_xyl_, the highest one among the overexpressed XK strains (Table 2; Figure 2b). The strain SBY140 was also the only one to show an improvement in μ_max_, 30%, compared with the parental strain (Figure 2a, Table 2). All these values were statically significant different (p-value ≤0.05) when compared with the values shown by the parental strain in the same conditions. On the other hand, this strain showed a lag phase of almost 30 hours while the parental strain had a lag phase of less than 20 hours (Figure 2d).

### 3.4. Characterization of double-transformant strain with heterologous expression of the genes ssXR + ciXK, and ssXR + ecXK

SBY137 showed improved growth characteristics compared to the parental strain (Table 2; Figure 2a-c), SBY140 showed the best improvement in terms of r_xyl_ and μ_max_ (Table 2; Figure 2a and b), and SBY141 improvement in r_xyl_ and short lag phase. Therefore, ssXR was inserted in SBY140 and SBY141 (Table 1), resulting in strains SBY165 and SBY166, respectively.

After the yeast transformation, three colonies were selected for each, and characterized using microplate cultivation, and based on the growth profile and μ_max_ calculation (Supplementary Figures 1 and 2) two candidates, SBY165 and SBY166 (Table 1), were selected for characterization in shake flasks. μ_max_, r_xyl_, Y_sx_ were calculated for the transformed strains (Table 2; Figure 2a-c). SBY166 showed an improvement of 118% in μ_max_ and 51% in r_xyl_. Different of others transformed strains, SBY166 was the only one that showed an improvement of 43% in Y_sx_ with a statically difference (p-value ≤0.05).

SBY165 showed an improvement of 118% in μ_max_ and 146% in r_xyl_ and (Figure 2a and 2b, Table 2), with a statically difference (p-value ≤0.05). No significant difference was seen in Y_sx_ compared to SBY92 (Figure 2c).

The parental strain produced 11.9 ± 1.3 mg.L^-1^ of carotenoids and the transformed strains SBY140, SBY141, SBY165, and SBY166 produced 1.6 ± 0.1, 4.7 ± 1.2, 7.7 ± 1.8, and 5.9 ± 0.1 respectively, while lipids content in biomass were 35.9% ± 0.2% for parental strain, 21.7% ± 0.4% for SBY140, 29.1± 3.0% for SBY141, 30.4% ± 1.4% for SBY165, and 32.6% ± 4.4% for SBY166 (Figure 2e and 2f). Interestingly, while the values for carotenoids and lipids produced by SBY165 were smaller than the parental strain, they were not statistically different. Similarly, the lipid content produced by SBY141 and SBY166 did not show a statistically significant difference compared to the parental strain.

## 4. Discussion

Xylose is an important substrate for bioprocesses that is currently underutilized by the industry. Our goal was to evaluate the possibility of using a heterologous NAD(P)H-dependent XR (SBY137), and by doing so, the xylose r_xyl_ in *R. toruloides* was improved 26% (Figure 2b, Table 2). Expressing the heterologous enzyme from *S. stipitis* may have improved the cofactor NADPH balance in the strain, and also, considering that the heterologous XR enzyme must have been expressed in active form, that an increase in the active XR pool resulted in higher reaction rates, as stated by enzyme kinetics laws. Since the r_xyl_ increased but μ_max_ remained the same, it is possible that some reaction downstream of XR could be limiting carbon flux through the central carbon metabolism.

Furthermore, the best results in terms of r_xyl_ and μ_max_ achieved in this work happened when both enzymes ssXR and ecXK were co-expressed. The r_xyl_ in SBY165 was improved in 146% (Figure 2B, Table 2) and the μ_max_ was improved in 118% (Figure 2a, Table 2). Also, when XK (native or heterologous) were overexpressed, improvements in r_xyl_ by 56% (SBY138), 46% (SBY139), and 66% (SBY140) (Figure 2b, Table 2) were observed suggesting that the low expression of the enzyme XK could be limiting the xylose metabolism in this yeast because once XK was overexpressed with a stronger promoter (pXYL), xylose consumption and growth were improved.

Ledesma-Amaro et al. (2016) [48] constructed a xylose-consuming *Y. lipolytica* by overexpressing ssXR, ssXDH, and the strain’s native XK, the latter being crucial for growth in xylose. The same has already been reported by Ho et al. (1998) [49] for *S. cerevisiae*. The authors observed that overexpression of native XK in addition to XR and XDH from *S. stipitis* showed an efficient xylose fermentation to ethanol and low concentration of xylitol, concluding that this strategy is better than the usual overexpression of only XR-XDH from *S. stipitis* in *S. cerevisiae*. Rodriguez et al. (2016) [50], after observing a weak expression of native XK in *Y. lipolytica* when it was cultivated in xylitol, suggested that the limited expression of XK could be the bottleneck for xylose and xylitol metabolism in this yeast. The same was observed by Wu et al. (2019) [51]. We have described a similar synergy between ssXR and ecXK in SBY165, since the native XK of *R. toruloides* is not expressed.

The r_xyl_ achieved in this work was 0.44 g_xyl_.g_dcw_^-1^.h^-1^ when ssXR and ecXK were jointly overexpressed (SBY165), followed by 0.30 ± 003 g_xyl_.g_dcw_^-1^.h^-1^ when only ecXK was expressed (SBY140). The former is one of the highest calculated r_xyl_ reported for *R. toruloides*, higher compared to the rates reported to *S. cerevisiae, Y. lipolytica* or *S. stipitis* [13,44,48]. Lopes et al. (2020) [53] cultivated *R. toruloides* in xylose (10 g.L^-1^) mineral medium and obtained an r_xyl_ of 0.52 g_xyl_.g_dcw_^-1^.h^-1^, however this was achieved by cultivating the yeast in bioreactors (turbidostat mode), which benefits cell growth by providing better mass transfer conditions of substrates, stable pH, and oxygen. Also, this value was achieved with a lower C/N (60 mol.mol^-1^) than our study, which has an inversely proportional effect on substrate uptake. When, in the same work, Lopes et al. (2020) [53] used a xylose C/N of 80, the r_xyl_ decreased to 0.39 g_xyl_.g_dcw_^-1^.h^-1^, a lower value than what we obtained in flasks. With all this considered, we may conclude that SBY165 has the potential of achieving even higher r_xyl_, if cultivated in conditions that allow doing so.

However, a significant decrease in lipids and carotenoid production was observed in SBY140 (Figure 2e, f). This could suggest a shortage of NADPH, necessary for lipid production. When ecXK was overexpressed in this strain with a stronger promoter, the NADPH may not have been regenerated more efficiently through the alternative pathway anymore, but likely through the oxidative part of the PPP, leading to less NADPH available for the lipids biosynthesis pathway, once the XR in this strain was the native one and NADPH-dependent. Nevertheless, when XR was added and both enzymes were overexpressed together (SBY165) the carotenoids increased almost 5-fold and total lipids increased by 40% compared with only ecXK being expressed (Figure 2f) and both values were not statically different when compared with the parental strain. This result could support the hypothesis that in the presence of XR from *S. stipitis*, the co-factor NADH was being used in the XR-XDH pathway instead NADPH, and more of this cofactor was available for the lipid biosynthesis pathway.

Clones harbouring the same heterologous enzyme showed statistically different specific growth rates compared to each other; some higher than SBY92, but some lower (Supplementary Figure 2). This is not expected when cultivating different clones obtained through site-directed integration if one considers that only one copy of the gene of interest was added to the genome. Therefore, we speculate it is very likely that additional random integration events have occurred for some or all the clones. In that case, the XR or XK cassettes may have integrated into different parts of the genome with effects that could have been either beneficial or deleterious to cell growth, which explains the variety of growth rates observed between clones from the same transformation.

It could be argued that the increase in growth could be related to these random integration events that caused a positive effect on it. However, an increase in growth rate is also an expected effect of overexpressing enzymes related to the uptake of carbon sources, which is the case of XR and XK, for the reason that they increase flux through the central carbon metabolism. In this case, the increase in growth rates would be coupled with an increase in xylose uptake rates. That was the case of the most of strains which were selected solely based on their μ_max_ in the microplate experiments, where the r_xyl_ was not calculated, but they all presented statistically higher r_xyl_ than their respective parental strains in the shake flask characterizations. For those cases, we conclude it is plausible to say that XR and/or XK were successfully overexpressed and resulted in the increase of the fermentation kinetic parameters. We believe the same conclusion applied to the SBY138 and SBY139 strains, even though it was not selected based on its μ_max_ but still showed an r_xyl_ 56% and 46% higher than its parental strain, respectively.

Another consequence of the occurrence of random integration events is that the expression of the inserted genes is affected by the chromosomal site where they were integrated in. This effect was already described in yeasts such as *S. cerevisiae* [54] and *Y. lipolytica* [55]. Zhang et al (2016) [54] reported 13-fold differences in gene expression over 1000 genomic loci in *S. cerevisiae*, a much higher magnitude than the increases in μ_max_ and r_xyl_ that were obtained in our work. Taking this into consideration limits our ability to compare the effects that each different XK had on growth and xylose uptake, for in the scope of this work we were not able to sort out the effect of the chromosomal position of our supposedly random integrated genes. Further studies need to be done using DNA sequencing to understand better how the integration in different positions on the chromosome can affect the physiology of the strain in this case.

## 5. Conclusions

The highest improvement in the specific-xylose consumption rate in SBY165 was observed when simultaneously overexpressing the heterologous XR and XK. This showed to be a good strategy for improving the xylose consumption rate in this yeast, making the strain produce statically the same amount of total lipids as the parental strain consuming the xylose 146% faster. Even with unknown gaps that still need to be solved about the xylose pathway in this yeast, the results presented in this work can support the previous hypothesis that the low expression of XK could be limiting its sugar consumption. Future studies could be carried out to increase even more the xylose consumption in *R. toruloides*, such as the overexpression of xylose transporter genes and adaptive evolution.

## Supplementary Materials

Figure S1: Growth profiles of selected colonies of transformant strains; Figure S2: Maximum specific growth rates (μ_max_) calculated for selected colonies of transformant strains; Table S1: A list of all primers used in this work.

## Author Contributions

PM, MJP, NB, and P-JL designed the experiments. PM, MJP, JSB, AT, and ET performed the experiments. PM, MJP, JSB, KH, NB, and P-JL analyzed the data. All authors wrote and revised the manuscript.

## Funding

This project has received funding from the European Union’s Horizon 2020 research and innovation program under Grant Agreement No 668997, and the Estonian Research Council grants PUT1488P and PRG1101. PM would additionally like to acknowledge, DORA Plus (nr. 2-1.17/TI/14).

## Data Availability Statement

The data presented in this study are available on request from the corresponding author.

## Acknowledgments

We thank Andreia Axelrud Nunes for help with HPLC analysis.

## Conflicts of Interest

The authors declare no conflict of interest. The funders had no role in the design of the study; in the collection, analyses, or interpretation of data; in the writing of the manuscript; or in the decision to publish the results.

## Notes

### Competing Interest Statement

The authors have declared no competing interest.

