## Supplementary Materials for "Improving xylose consumption in *Rhodotorula toruloides* through heterologous expression of xylose reductase and xylulokinase"

**Table S1.** A list of all primers used in this work.

| **Sequence (5’ – 3’)** | **Purpose** |
| --- | --- |
| tctctgggtctcg*ggag*ttgacgcgcgactttgcg | Amplification of insUP PEX10 + NAT + tNOS + pXYL for GGA assembly |
| atgc*ggtctc*acattcgacatggcgtgtattctg |  |
| gcat*ggtctc*atctacgttcaaacatttggcaataaagtttc | Amplification of tNOS + insD PEX10 for GGA assembly |
| tctctgggtctcc*atgg*agtcgttcgaatggttct |  |
| tctctgggtctcg*ggag*ttgacgcgcgactttgcg | Amplification of upstream of PEX10 gene |
| tctctgggtctcg*acct*gtcggctgctgtgttg |  |
| tctctgggtctcc*gtgt*ttgctgaacgtgaagc | Amplification of downstream of PEX10 gene |
| tctctgggtctcc*atgg*agtcgttcgaatggttct |  |
| *tctctgggtctcg*ggagctgaccagtccaaagggaagaccag | Amplification of upstream of PEX11 gene |
| atgc*ggtctc*aaccttcctggactaaccatggctggttg |  |
| *tctctgggtctcg*gtcagcaaggtaaagggggcgtctgcgtc | Amplification of downstream of PEX11 gene |
| atgc*ggtctc*aatggatcggctacctcgtcaagctcatcc |  |
| gcat*ggtctca*aatgccctcgatcaagctcaactcg | Amplification of ssXR gene for GGA assembly |
| atgc*ggtctca*tagattagacgaagatggggatcttgtccc |  |
| acggggtctcc*aatg*atgacttataaggaagagactggc | Amplification of ciXK gene for GGA assembly |
| acggggtctcg*taga*ctagtcgtgcttgagctg |  |
| acggggtctcc*ggag*atgtacatcggcatcgac | Amplification of ecXK gene for GGA assembly |
| acggggtctcg*atgg*ctacgccataagtgggag |  |
| acggggtctccaatgatgaccacgaccccattc | Amplification of ssXK gene for GGA assembly |
| acggggtctcgtagactagtgtttgagctccgactcc |  |
| gcatggtctcaccacatgaagaaacctgagcttaccgc | Amplification of HYG gene for PCR verification |
| atgcggtctcaatacctactctttagcgcgaggtcg |  |
| atgggtccttcaccaccg | Amplification of NAT gene for PCR verification |
| gagatgaccacgaagcccg |  |

**Table S1.** A list of all primers used in this work (continuation).

| **Sequence (5’ – 3’)** | **Purpose** |
| --- | --- |
| gatgtgggtctcc*aggt*tgtccgtattctacatcga | Amplification of promoter pXYL for PCR verification |
| gatgtgggtctcc*tgat*cgacatggcgtgtattctg |  |
| gatgtgggtctcc*aggt*cggctgaggcttccccg | Amplification of promoter pADH2 for PCR verification |
| gatgtgggtctcc*tgat*tgtgactgtcggagacgtggca |  |
| ctcctcctcgacctcctccg | Amplification of ssXR gene for PCR verification |
| gggacggtgttcgacttggg |  |
| cgagggcccactaccgcttc | Amplification of ciXK gene for PCR verification |
| agcgctccgttgcagtagca |  |
| acgccacacaacaaccccca | Amplification of ecXK gene for PCR verification |
| ttcgcgtcgtggctggtagg |  |
| tcttcaaccccgtcctcg | Amplification of rtXK gene for PCR verification |
| cctcatcttcccgatcttggt |  |
| cataactctaatgtcatttaacgttcaaacatttggcaataa | Amplification of terminator tNOS gene for PCR verification |
| gcgtcggggaagcctcagcccccgatctagtaacatagatga |  |

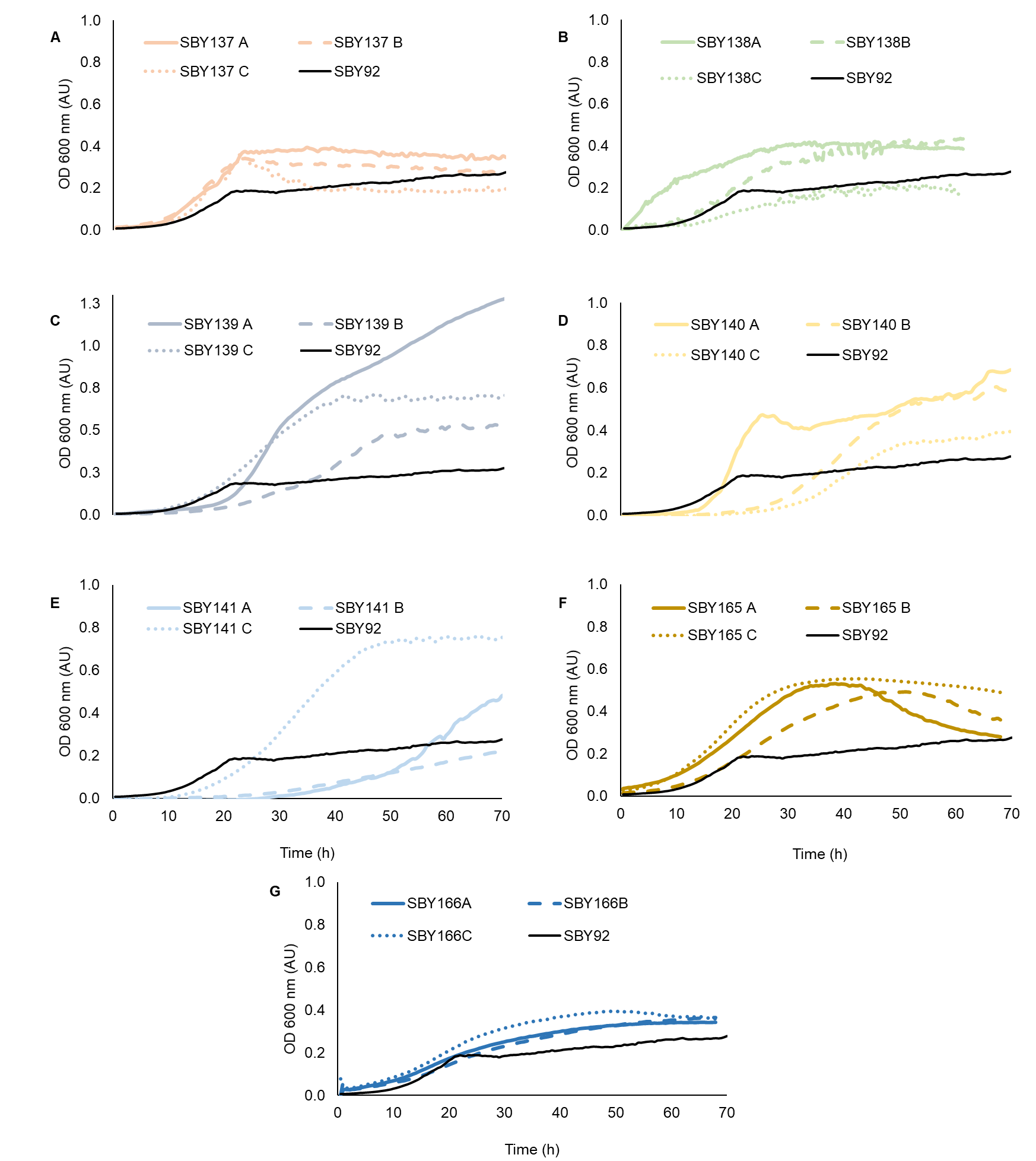

**Figure S1.** Growth profiles of selected colonies of transformant strains. (**a**) selected colonies of transformant strain SBY137; (**b**) selected colonies of transformant strain SBY138; (**c**) selected colonies of transformant strain SBY139; (**d**) selected colonies of transformant strain SBY140; (**e**) selected colonies of transformant strain SBY141; (**f**) selected colonies of transformant strain SBY165; (**g**) selected colonies of transformant strain SBY166. Characterization was carried out on microplate cultivation using mineral medium and 1 g/L of xylose. The OD was acquired every 30 min and the curves represent the average of at least 3 replicates.

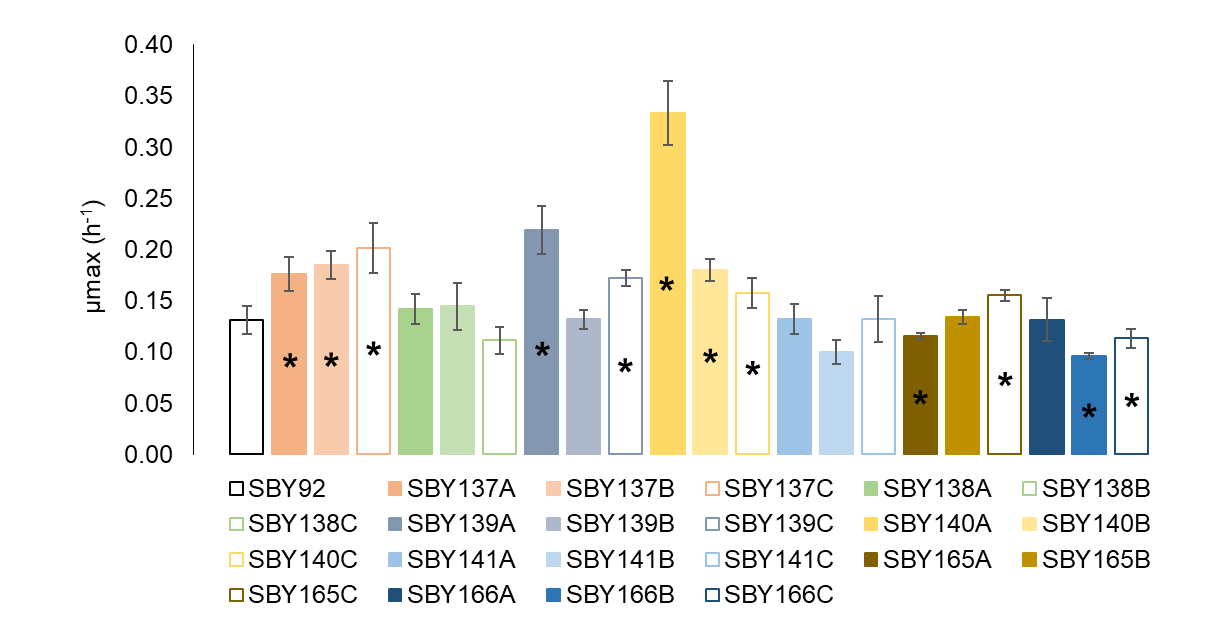
**Figure S2.** Maximum specific growth rates (µ_max_) calculated for selected colonies of transformant strains. Strains were characterizated on microplate cultivation using mineral medium and 1 g/L of xylose. All µmax were calculated using the growth profile and represent the average of at least 3 replicates. A, B, C are different colonies characterized of each transformation. *values are statistically significant different with p-value ≤ 0.05 when compare with the µmax obtained with the parental strain (SBY92).
